# The Sentieon Genomics Tools - A fast and accurate solution to variant calling from next-generation sequence data

**DOI:** 10.1101/115717

**Authors:** Donald Freed, Rafael Aldana, Jessica A. Weber, Jeremy S. Edwards

## Abstract

In the past six years worldwide capacity for human genome sequencing has grown by more than five orders of magnitude, with costs falling by nearly two orders of magnitude over the same period [1], [2]. The rapid expansion in the production of next-generation sequence data and the use of these data in a wide range of new applications has created a need for improved computational tools for data processing. The Sentieon Genomics tools provide an optimized reimplementation of the most accurate pipelines for calling variants from next-generation sequence data, resulting in more than a 10-fold increase in processing speed while providing identical results to best practices pipelines. Here we demonstrate the consistency and improved performance of Sentieon’s tools relative to BWA, GATK, MuTect, and MuTect2 through analysis of publicly available human exome, low-coverage genome, and high-depth genome sequence data.

## Introduction

The cost of sequencing a human genome has fallen rapidly over the last decade and is now near $1000 for a single high-depth human genome, with annual increases in both sequencing efficiency in Gb per dollar and worldwide sequencing capacity consistently surpassing Moore’s Law [1], [2]. Importantly, data quality has also improved due to advances in the underlying technology and chemistry of the sequencing machines. Read lengths of the Illumina’s flagship X10 sequencer are now 150bp and data quality remains high across the entire read [3].

As the amount of available sequence data increases, efficient and accurate data analysis is becoming increasingly important. Next-generation sequence data are frequently being used to help inform economic and clinical decisions through applications such as non-invasive prenatal testing [4], personalized therapy including cancer immunotherapy [5]–[10], genetic diagnosis [11], disease gene discovery [12], [13], discovery of contributory mutations in complex disease [14], and discovery of important genetic traits in agriculture [15]. Due to the increased reliance upon next-generation sequence data for informing these decisions, the pipelines for analyzing these data are understandably under increasing regulatory scrutiny [16].

While early human genome projects necessarily relied upon whole-genome *de novo* assembly, the short-read data produced by second-generation sequencers are not readily amenable to this method of assembly. As a result, many common data analysis pipelines involve mapping sequence reads to a reference genome and identifying single nucleotide variants (SNVs) and other variants relative to the reference [17]–[20]. Initially these methods relied upon Bayesian approaches to evaluate the likelihoods of mutations occurring at single-base pairs. However, newer haplotype-based approaches have improved accuracy and have become the industry standard [21]–[23].

Two of the most popular tools for variant detection are the GATK and MuTect [23]–[25]. The publications describing these tools have been cited nearly 10,000 times and both have been used in many high-profile research projects [14], [26], [27]. Part of the reason for their wide adoption is their high accuracy; these tools perform well in many benchmarks including the precisionFDA and ICGC-TCGA DREAM challenges [28], [29]. However, these tools do have drawbacks. The improvements that produce higher accuracy have also resulted in long runtimes and high memory usage for some tasks. Downsampling is implemented in some algorithms to help mitigate these issues, but is not an ideal solution as it discards valuable data, leads to increased run-to-run variation, and possibly results in erroneous or missing variant calls. In some steps, intermediate file merging is also necessary to reduce computational load.

Due to the projected increase in the amount of available next-generation sequence data and the use of these data for increasingly important applications, there is a pressing need for new and improved software solutions for next-generation sequence data analysis. We have developed the Sentieon Genomics tools to address this need. The Sentieon tools provide a complete rewrite of the mathematical models of the Best Practices GATK, Picard, MuTect, and MuTect 2 in the Sentieon DNAseq, TNseq and TNHaplotyper pipelines with a focus on computational efficiency, accuracy, and consistency.

## Materials and Methods

Benchmarking comparisons were completed using BWA 0.7.12-r1039 [30], [31], Picard tools 1.112, GATK 3.5 [23], [24], SAMtools 1.2 [32], MuTect 1.1.5 [25], and version 201611 of the Sentieon Genomics tools. All commands were run on a 32 core 2.4 GHz Intel Xeon server with 64 GB memory and 2TB dual stripped SSDs for intermediate file storage. The server was dedicated to benchmarking and had no other running jobs. All samples used in benchmarking are publicly available and are listed in Supplementary Table 1. Data from multiple lanes that were obtained from the Baylor Human Genome Sequencing Center were concatenated into a single file before analysis. Samples HG001 and HG002 were obtained from the precisionFDA platform.

For all samples, sequence reads were aligned to the Human reference genome (UCSC hg19) with BWA-MEM followed by sorting and indexing using samtools or the Sentieon utility. Alignment summary, GC bias, base quality by sequencing cycle, base quality score distribution, and insert size metrics were collected and duplicate reads were removed with either Picard tools or the Sentieon driver. Indels were realigned and base quality was recalibrated using either the GATK or the Sentieon driver. For tumor-normal paired samples, joint indel realignment was performed using the GATK or the Sentieon driver. For germline samples, variants were called using the GATK UnifiedGenotyper and the GATK HaplotypeCaller or Sentieon DNAseq Haplotyper and Genotyper algorithms while variants were called in paired samples using MuTect and MuTect2 or Sentieon TNseq TNsnv and TNHaplotyper algorithms.

For the benchmarking of the joint calling, gVCF files were generated using the Sentieon pipeline described in Supplementary Appendix 1 with the --emit_mode gvcf option added during variant calling. gVCF files were then genotyped by the GATK GenotypeGVCFs and Sentieon DNAseq GVCFtyper across chromosome 1. Genotyping was not performed across the entire genome as the estimated runtime for the whole-genome analysis with the GATK was over two weeks. Since Sentieon’s GVCFtyper does not drop alternate alleles, GenotypeGVCFs was run with the option-maxAltAlleles 100 to provide comparable results.

RTG Tools’ vcfeval 3.5.1 was used to compare variants called by Sentieon to variants called by the other tools. Detailed commands used during the data processing are listed in Supplementary Appendix 1.

## Results

### The Sentieon Genomics Pipeline Tools

The Sentieon Genomics pipeline provides a suite of tools for secondary analysis of next-generation sequence data. Currently supported pipelines are composed of optimized implementations of the mathematical models of the most accurate variant calling pipelines. Improvements in performance are achieved through optimization of the algorithms and improved resource management. The tools run on both Linux (RedHat/CentOS, Debian, OpenSUSE and Ubuntu) and OS X distributions and require no specialized hardware, additional libraries or complex installation procedure. Sentieon’s variant calling tools do not perform data down-sampling and are deterministic, providing perfect run-to-run consistency [28].

### The Sentieon Tools Provide a 10-fold Improvement in Runtime Over the GATK, MuTect and MuTect2

To test the performance of the Sentieon Genomics pipeline tools, we evaluated the runtime of Sentieon’s tools and the GATK, MuTect, and MuTect2 in a consistent computing environment using publicly available data (a complete set of samples is listed in Supplementary Table 1). Sentieon’s tools provide near optimal parallelization with built-in multithreading functionality. Unfortunately, MuTect is not multithreaded and the built-in multithreading of the GATK and MuTect2 are known to be suboptimal. While advanced users may use sophisticated parallelization methods to improve performance (such as performing operations on small intervals and concatenating the results), optimization, validation and testing of these methods require expertise, expensive human capital investment, and these methods vary in their effectiveness from user-to-user. Due to these costs, only built-in parallelization methods are tested here. Multithreaded processing of the Sentieon algorithms was accomplished using the “-t” option while the GATK, and MuTect2 were parallelized with the “-nct” option. The full set of commands used in all analyses is shown in Supplementary Appendix 1.

Prior to variant calling, sequence data were aligned to the human reference genome (UCSC hg19) using BWA-MEM. BWA is one of the most popular aligners for alignment of next-generation sequence reads given its accuracy and ability to produce correct alignments at structural variant breakpoints. Sentieon provides an optimized implementation of BWA resulting in an average 1.9x speedup (range 1.0 to 3.9x, Figure 1, Supplementary Table 2), while producing identical alignments. Sentieon DNAseq and the GATK Best Practices Pipeline were run on seven whole-exome, 78 low-coverage, and two whole-genome samples. Among all samples, the Sentieon DNAseq pipeline resulted in an average 36x (range 16 to 55x) improvement in runtime relative to the GATK Best Practices pipeline (Figure 2, Supplementary Table 2, Supplementary Table 3). The performance improvements were most notable for the indel realignment, base-quality score recalibration, and HaplotypeCaller variant calling stages where Sentieon’s tools improved runtimes by an average of 56x, 46x, and 30x, respectively.

**Figure 1.**
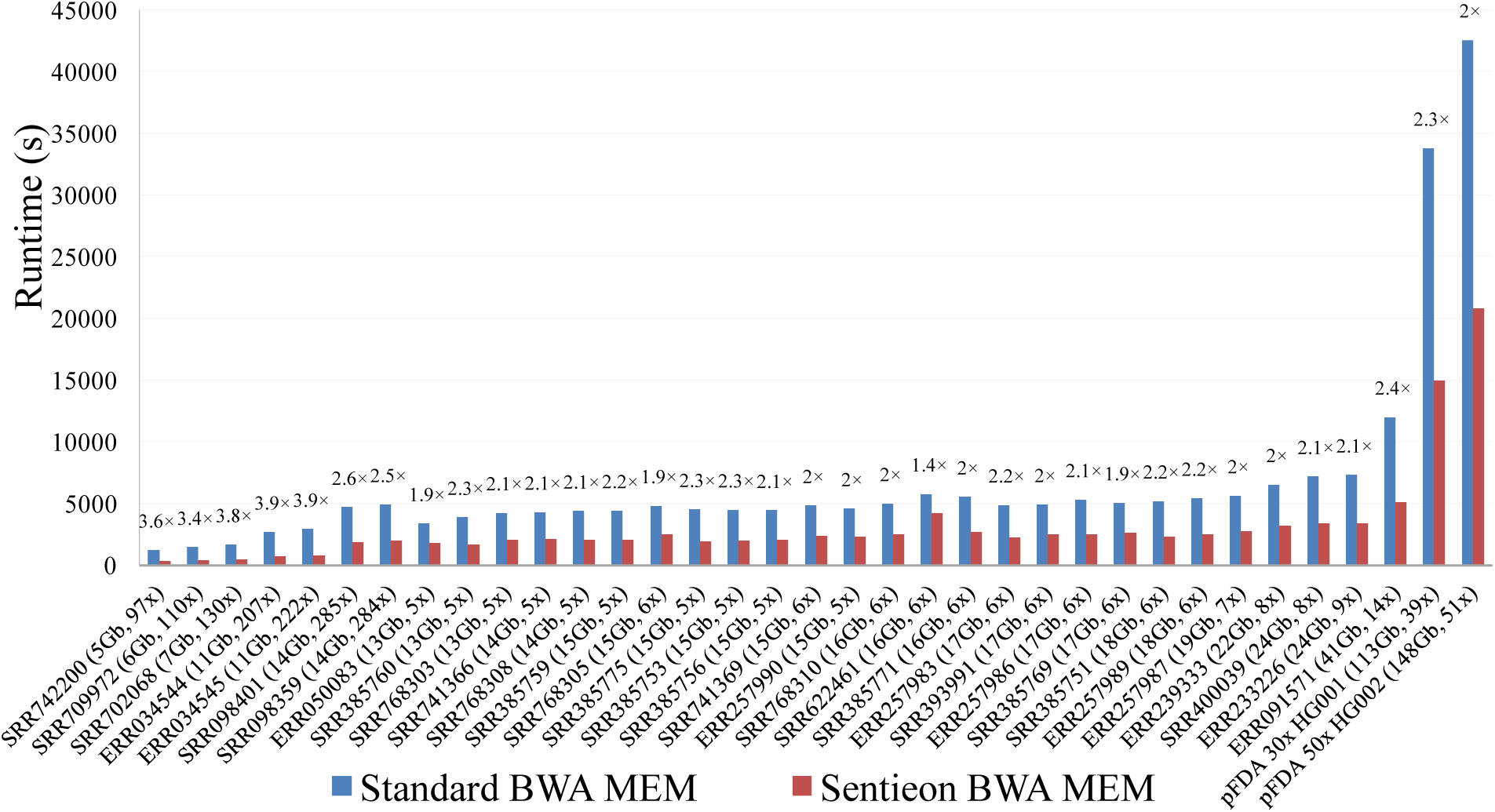
Standard BWA-MEM and Sentieon BWA-MEM runtime comparison. Runtimes of the standard BWA with SAMtools sort and Sentieon BWA and sort on whole-exome and low-coverage whole-genome and high-coverage whole-genome samples. Labels indicate fold improvement in runtime provided by the Sentieon implementation.

**Figure 2.**
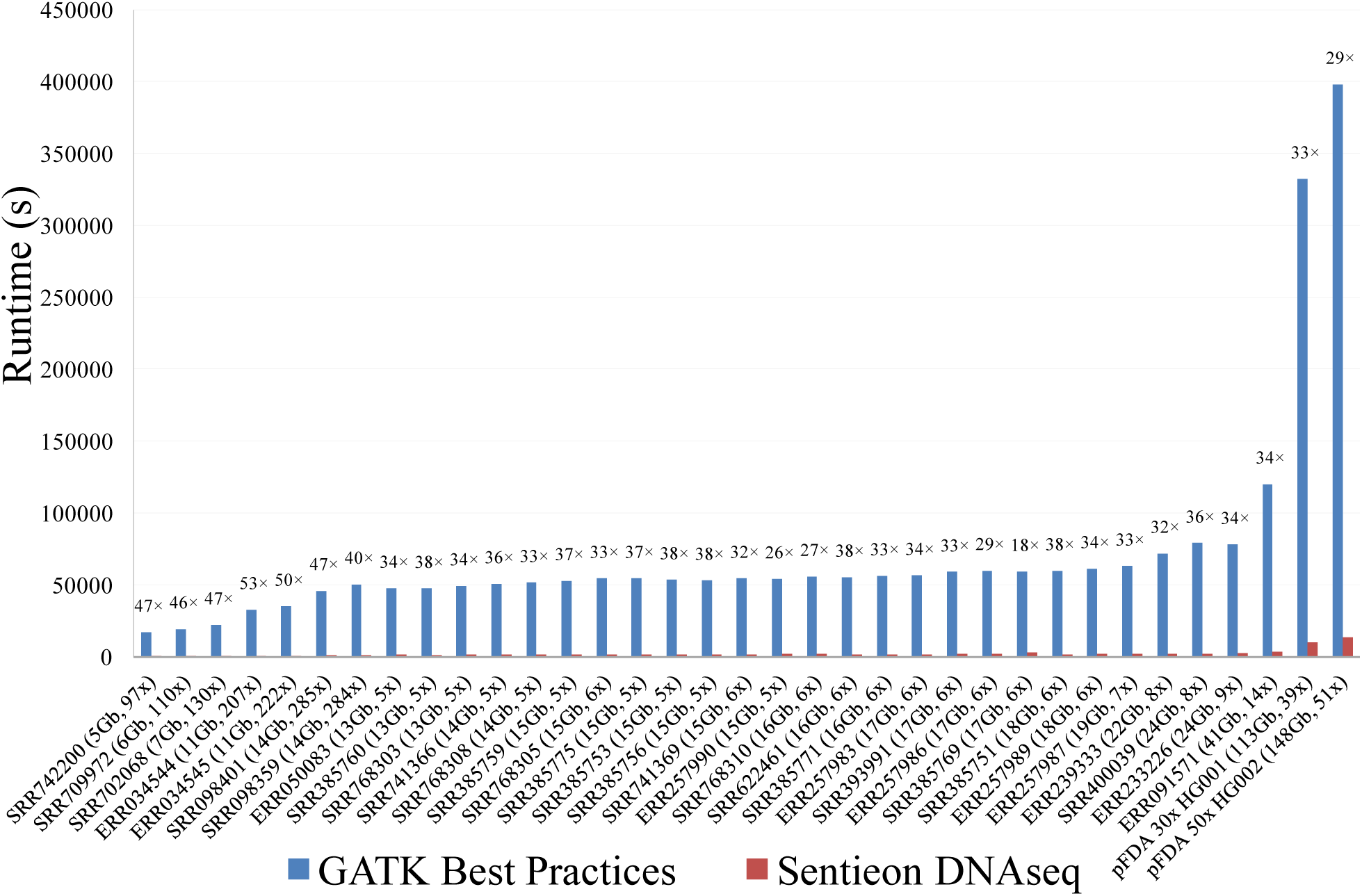
DNAseq pipeline runtime comparison. Runtimes of the Sentieon DNAseq and GATK Best Practices pipelines on whole-exome, low-coverage whole-genome, and high-coverage whole-genome samples for the metrics calculation through variant calling stages. Samples were sorted by their total number of sequenced bases. Labels indicate the fold improvement in runtime provided by the Sentieon tools over the GATK. The runtime improvement of Sentieon DNAseq over GATK ranges from 18-53x.

To better understand the performance differences between the GATK and Sentieon tools for joint genotyping, we ran the GATK GenotypeGVCFs and Sentieon DNAseq GVCFtyper on chromosome 1 of 74 gVCF files from the 1000 Genomes Project preprocessed with the Sentieon DNAseq pipeline. Sentieon’s tools provided a 183x improvement in runtime relative to the GATK for joint genotyping (216 and 39,555 seconds for Sentieon and the GATK, respectively).

We also measured the performance of the Sentieon TNseq TNsnv and TNhaplotyper pipelines relative to MuTect and MuTect2 with three tumor-normal paired samples. Sentieon’s tools resulted in an average speedup of 19x (range 10x to 24x) relative to MuTect2 and an average speedup of 42x (range 38 to 48x) relative to the single-threaded MuTect (Figure 3, Supplementary Table 4). Overall, the Sentieon tools provide an average 19x to 42x speedup over earlier implementations of the variant calling pipelines in the GATK, MuTect and MuTect2.

**Figure 3.**
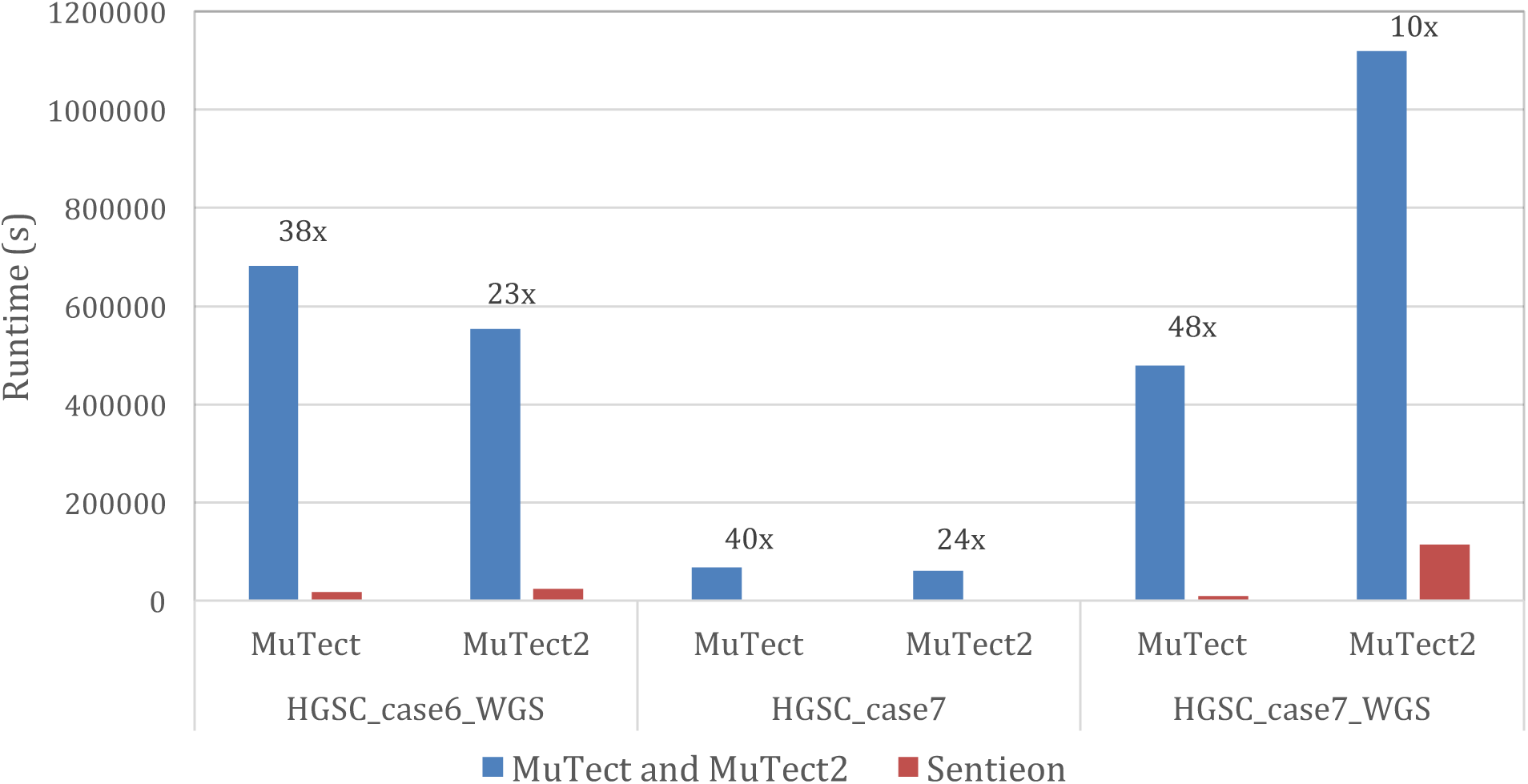
TNseq and TNhaplotyper runtime comparison. Runtimes of the Sentieon TNseq TNsnv and TNhaplotyper pipelines relative to MuTect and MuTect2 on whole-exome and whole-genome samples. Stages included in the plot are metrics calculation through variant calling. Labels indicate fold improvement in runtime provided by the Sentieon tools relative to MuTect and MuTect2.

### The Sentieon Tools Produce Results Consistent with the GATK, MuTect and MuTect2

Using the VCF files produced in the runtime benchmark, we set out to evaluate the consistency of Sentieon’s tools with the GATK, MuTect, and MuTect2 using RTG Tools vcfeval. When interpreting the results of these experiments, it is important to note the run-to-run variation of the tools. While the Sentieon tools are deterministic, repeated runs of multithreaded GATK and MuTect2 produce slightly different results, limiting the consistency that can be achieved between Sentieon and these other tools [33]. MuTect downsamples sequence data deterministically while GATK and MuTect2 downsample sequence data randomly when running with multiple threads. However, even deterministic downsampling may cause additional variants to be called or lost relative to an identical implementation without downsampling. Here we call the results identical if the measured difference between the tools is on the order of the intrinsic run-to-run variation of the GATK observed by Weber *et al.* [33]. The Sentieon DNAseq pipeline produced results identical to the GATK best practices pipeline (average F-score 0.9996; range 0.9974 to 1.0000; Figure 4; Supplementary Table 5). In the joint genotyping comparison, the Sentieon tools produced results identical to the GATK (Supplementary Table 6). While the Sentieon TNseq TNsnv and TNhaplotyper pipelines produced results identical to MuTect and MuTect2 (Average F-score 0.9994 and 0.9920 for MuTect and MuTect2, respectively; Supplementary Table 7). These results indicate that the Sentieon DNAseq and TNseq pipelines provide a drop-in replacement for the best-practices pipelines with results within the run-to-run variation of the tools.

**Figure 4.**
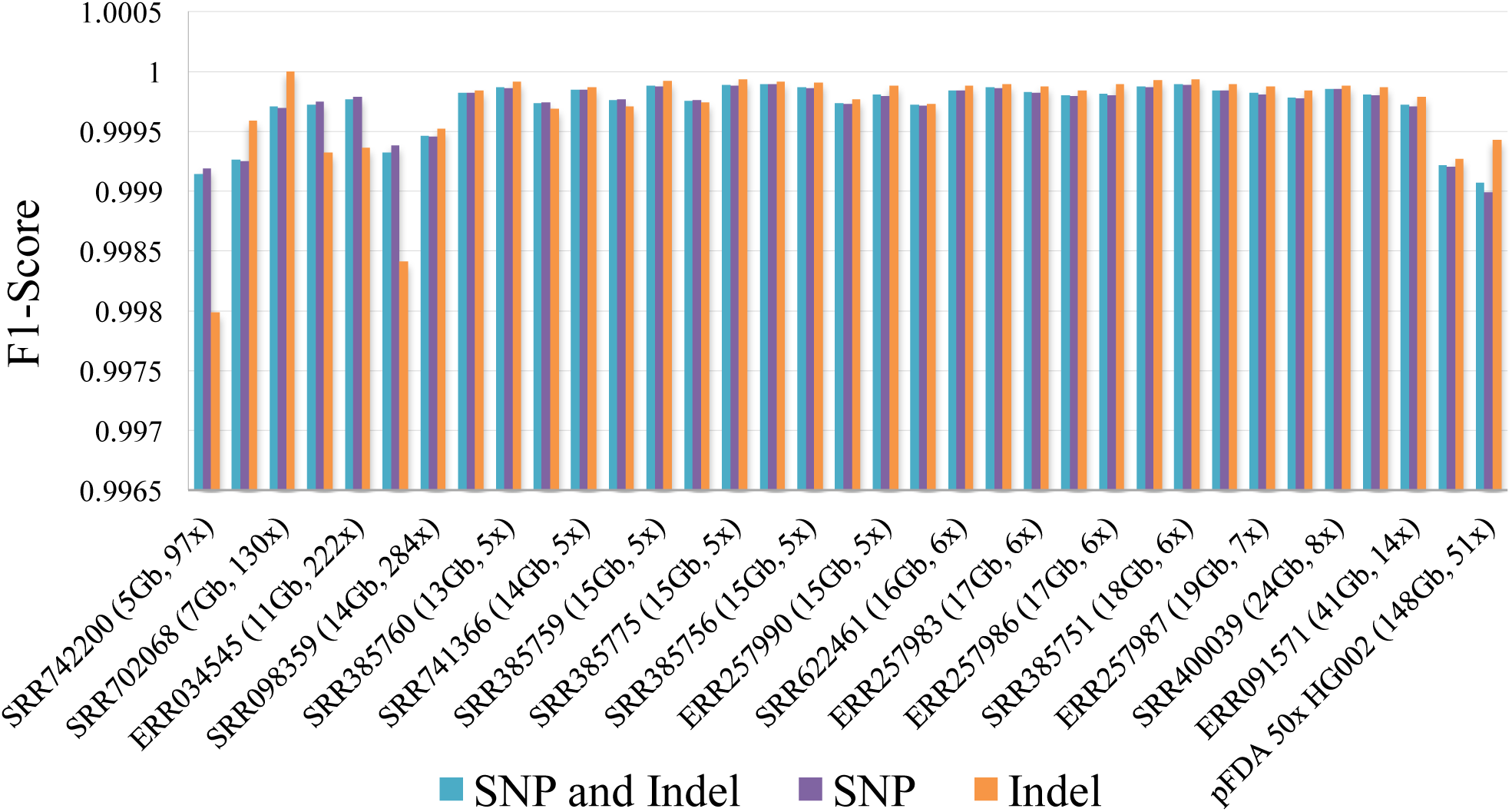
Consistency of Sentieon DNAseq and the GATK Best Practices. F1-Score between the Sentieon DNAseq and GATK Best Practices variant calls is shown for combined SNPs and Indels and each separately. F1-score was measured using RTG tools vcfeval.

## Discussion

In this manuscript, we present the Sentieon DNAseq and TNseq pipelines. The Sentieon tools provide computationally efficient, multithreaded, deterministic variant calling from germline samples and tumor-normal pairs with results identical to the GATK, MuTect and MuTect2 while providing a greater than 10-fold improvement in total runtime. As it provides identical results, the Sentieon tools can function as a drop-in replacement for the GATK, MuTect or MuTect2 resulting in cost-savings for researchers and clinicians. The lack of down-sampling enables high-depth sequencing for increased accuracy, the robust implementation enables joint genotyping of 100,000s of files simultaneously without intermediate file merging, and the algorithmic determinism provides the consistency required for medical applications. As next-generation sequence data is produced at an increasing rate and more frequently finds use in important economic and clinical applications, the Sentieon Genomics pipeline tools provide a means to process this data with accuracy, efficiency and consistency.

## Acknowledgements

We would like to thank the individuals who consented to making their genetic data publicly available.

We would like to thank the 1000 Genomes Project and EMBL-EBI for making the 1000 Genomes data publicly available. We thank the Texas Cancer Research Biobank and Baylor College of Medicine Human Genome Sequencing Center for providing the paired tumor-normal samples.

## References

[1] Z. D. Stephens et al., “Big Data: Astronomical or Genomical?,” PLoS Biol., vol. 13, no. 7, p. e1002195, Jul. 2015.

[2] NHGRI, “The Cost of Sequencing a Human Genome.” [Online]. Available: https://www.genome.gov/sequencingcosts/. [Accessed: 01-Feb-2017].

[3] Illumina, “HiSeq X Series of Sequencing Systems.” [Online]. Available: https://www.illumina.com/content/dam/illumina-marketing/documents/products/datasheets/datasheet-hiseq-x-ten.pdf. [Accessed: 30-Jan-2017].

[4] D. W. Bianchi et al., “DNA Sequencing versus Standard Prenatal Aneuploidy Screening,” N. Engl. J. Med., vol. 370, no. 9, pp. 799–808, Feb. 2014.

[5] E. A. Ashley, “Towards precision medicine,” Nat. Rev. Genet., vol. 17, no. 9, pp. 507–522, Aug. 2016.

[6] N. Hacohen, E. F. Fritsch, T. A. Carter, E. S. Lander, and C. J. Wu, “Getting Personal with Neoantigen-Based Therapeutic Cancer Vaccines,” Cancer Immunol. Res., vol. 1, no. 1, pp. 11–15, Jul. 2013.

[7] J. P. Finnigan, A. Rubinsteyn, J. Hammerbacher, and N. Bhardwaj, “Mutation-Derived Tumor Antigens: Novel Targets in Cancer Immunotherapy,” Oncol. Williston Park N, vol. 29, no. 12, Dec. 2015.

[8] M. A. Hamburg and F. S. Collins, “The Path to Personalized Medicine,” N. Engl. J. Med., vol. 363, no. 4, pp. 301–304, Jul. 2010.

[9] T. N. Schumacher and R. D. Schreiber, “Neoantigens in cancer immunotherapy,” Science, vol. 348, no. 6230, pp. 69–74, Apr. 2015.

[10] S. J. Aronson and H. L. Rehm, “Building the foundation for genomics in precision medicine,” Nature, vol. 526, no. 7573, pp. 336–342, Oct. 2015.

[11] J. Thevenon et al., “Diagnostic odyssey in severe neurodevelopmental disorders: toward clinical whole-exome sequencing as a first-line diagnostic test: Diagnostic odyssey in severe neurodevelopmental disorders,” Clin. Genet., vol. 89, no. 6, pp. 700–707, Jun. 2016.

[12] S. B. Ng et al., “Exome sequencing identifies the cause of a mendelian disorder,” Nat. Genet., vol. 42, no. 1, pp. 30–35, Jan. 2010.

[13] K. M. Boycott, M. R. Vanstone, D. E. Bulman, and A. E. MacKenzie, “Rare-disease genetics in the era of next-generation sequencing: discovery to translation,” Nat. Rev. Genet., vol. 14, no. 10, pp. 681–691, Sep. 2013.

[14] I. Iossifov et al., “The contribution of de novo coding mutations to autism spectrum disorder,” Nature, vol. 515, no. 7526, pp. 216–221, Nov. 2014.

[15] H. D. Daetwyler et al., “Whole-genome sequencing of 234 bulls facilitates mapping of monogenic and complex traits in cattle,” Nat. Genet., vol. 46, no. 8, pp. 858–865, Jul. 2014.

[16] R. B. Altman et al., “A research roadmap for next-generation sequencing informatics,” Sci. Transl. Med., vol. 8, no. 335, p. 335ps10, Apr. 2016.

[17] H. Li, J. Ruan, and R. Durbin, “Mapping short DNA sequencing reads and calling variants using mapping quality scores,” Genome Res., vol. 18, no. 11, pp. 1851–1858, Nov. 2008.

[18] R. Li et al., “SNP detection for massively parallel whole-genome resequencing,” Genome Res., vol. 19, no. 6, pp. 1124–1132, Jun. 2009.

[19] J. Wang et al., “The diploid genome sequence of an Asian individual,” Nature, vol. 456, no. 7218, pp. 60–65, Nov. 2008.

[20] T. J. Ley et al., “DNA sequencing of a cytogenetically normal acute myeloid leukaemia genome,” Nature, vol. 456, no. 7218, pp. 66–72, Nov. 2008.

[21] E. Garrison and G. Marth, “Haplotype-based variant detection from short-read sequencing.”

[22] A. Rimmer et al., “Integrating mapping, assembly and haplotype-based approaches for calling variants in clinical sequencing applications,” Nat. Genet., vol. 46, no. 8, pp. 912–918, Aug. 2014.

[23] M. A. DePristo et al., “A framework for variation discovery and genotyping using next-generation DNA sequencing data,” Nat. Genet., vol. 43, no. 5, pp. 491–498, May 2011.

[24] A. McKenna et al., “The Genome Analysis Toolkit: a MapReduce framework for analyzing next-generation DNA sequencing data,” Genome Res., vol. 20, no. 9, pp. 1297– 1303, Sep. 2010.

[25] K. Cibulskis et al., “Sensitive detection of somatic point mutations in impure and heterogeneous cancer samples,” Nat. Biotechnol., vol. 31, no. 3, pp. 213–219, Mar. 2013.

[26] A. Auton et al., “A global reference for human genetic variation,” Nature, vol. 526, no. 7571, pp. 68–74, Sep. 2015.

[27] N. Stransky et al., “The Mutational Landscape of Head and Neck Squamous Cell Carcinoma,” Science, vol. 333, no. 6046, pp. 1157–1160, Aug. 2011.

[28] “precisionFDA.” [Online]. Available: https://precision.fda.gov/. [Accessed: 27-Jan-2017].

[29] A. D. Ewing et al., “Combining tumor genome simulation with crowdsourcing to benchmark somatic single-nucleotide-variant detection,” Nat. Methods, vol. 12, no. 7, pp. 623–630, May 2015.

[30] H. Li and R. Durbin, “Fast and accurate short read alignment with Burrows-Wheeler transform,” Bioinformatics, vol. 25, no. 14, pp. 1754–1760, Jul. 2009.

[31] H. Li, “Aligning sequence reads, clone sequences and assembly contigs with BWA-MEM,” ArXiv Prepr. ArXiv13033997, 2013.

[32] H. Li et al., “The Sequence Alignment/Map format and SAMtools,” Bioinformatics, vol. 25, no. 16, pp. 2078–2079, Aug. 2009.

[33] Jessica A. Weber, Rafael Aldana, Brendan D. Gallagher, and Jeremy S. Edwards, “Sentieon DNA pipeline for variant detection - Software-only solution, over 20× faster than GATK 3.3 with identical results,” 2016.

